# Phylogeography and Evolutionary analysis of African Rotavirus A genotype G12 reveals district genetic diversification within Lineage III

**DOI:** 10.1101/591297

**Authors:** Babatunde Olanrewaju Motayo, Olukunle Oluwasemowo, Babatunde Adebiyi Olusola, Adewale Victor Opayele, Adedayo Omotayo Faneye

**Author notes:** Corresponding Author BABATUNDE OLANREWAJU MOTAYO, Department of Virology, College of Medicine, University of Ibadan, Nigeria. Postal address: Pathology department, Federal Medical Centre, P.M.B. 3031 Sapon, Abeokuta, Nigeria.

## Abstract

Rotavirus genotype G12 has become one of the most prevalent genotypes of rotavirus in Africa. To understand the drivers for its genetic diversity we investigated the Bayesian phylogeney, evolution and population demography of the genotype G12 Africa. Rotavirus genotype G12, VP7 sequences were downloaded and aligned from twelve African countries (n=96). Phylogenetic analysis, Evolutionary analysis and Bayesian Phylogeography was carried out, using MEGA Vs 6, BEAST, and SPREAD3. Phylogeny showed that all the African sequences fell into lineage III diversifying into two major clades. The evolutionary rate was 1.678×10^-3^ (95%HPD, 1.201×10^-3^ -2.198×10^-3^) substitutions/ site/ year. The MCC tree topology clustered into three lineages (II, III, IV), African strains diversified into three clusters within lineage III. South Africa was the epicentre of viral dispersal. This study shows the potential for genetic diversification of Rotavirus G12 in Africa, continuous molecular surveillance across Africa is recommended to help control effort.

**Highlights:** Our study revealed that African G12 rotaviruses have diversified into 3 clades within their parental lineage III based on geographic boundaries.

Nigeria was identified Nigeria as country of origin, while South Africa served as the epicentre of dispersal of the genotype across Africa.

We also discovered that they have a constant demographic profile. Our findings reveal the potential for rapid genetic diversity of Rotavirus G12 and highlight the importance of molecular surveillance in Rotavirus control effort.

## 1. Introduction

Group A rotavirus (RVA) has been established to be the main agent responsible for acute gastroenteritis (AGE) among children and infants worldwide. In 2016, it was reported to be responsible for about 128,000 deaths with over 2/3 of cases occurring in sub-Saharan Africa [Troeger et al, 2018]. There are two life attenuated rotavirus vaccines, Rotarix and Rotateq, which have been licensed for use in many countries after large phase 3 clinical trials were conducted in 2006 [Ruiz-Palacious et al, 2006, Vesikari et al, 2006]. In 2009 the world health organisation WHO, recommended the global use of the 2 live attenuated vaccines Rotarix and Rotateq. In Africa, several countries have included rotavirus vaccination in their EPI programs [Jonestella et al, 2017].

Rotavirus belongs to the virus family Reoviridae, it is a non-enveloped and has an icosahedral nueleocapsid structure, enclosing a double stranded (ds) RNA genome segmented into 11 compartments. The rotavirus RNA genome codes for six structural proteins, (VP1 to VP4, VP6 and VP7) and five/ six nonstructural proteins (NSP1 to NSP5/6)[Estes and Greenberg, 2013]. There areat least 10 distinct species/groups (A-I, J), differentiated by their VP6 antigenic properties [Banyai et al, 2017]. There are 32 G (VP7) genotypes and 47 P (VP4) genotypes identified through molecular epidemiologyhttps://rega.kuleuven.be/cev/viralmetagenomics/virus-classification/7th-RCWG-meeting.updateof the Rega Institute, KU Leuven, Belgium.

Molecular epidemiology has identified the widespread circulation of various genotypes of rotavirus in Africa, the globally prevalent genotypes G1P[8], G3P[4] have been reported in various African countries [Simwaka et al, 2018; Mwenda et al, 2018; Lartey et al, 2018; Moure et al, 2018,11Motayo et al, 2018]. However these globally prevalent genotypes are being replaced by more recent genotypes such as G9P[8], G8P[4], G12P[6], G12P[8], [12 Japhet et al, 2012]. There have also been recent reports of widespread outbreaks in parts of West Africa such as Nigeria attributed to RVA G12 strains [12 Japhet et al, 2012; Ianiro et al, 2015; Japhet et al, 2018]. Report has also suggested the rapid evolution of RVA genotype G12 as a panacea for its global spread [Matthijnssens et al, 2011]. The rapid emergence of this genotype in Africa has not been well understood, however mechanisms such as interspecies recombinations, RNA polymerase infidelity and accumulated point mutations have been attributed to rotavirus evolution [Estes and Greenberg 2013; Rahman et al, 2007; Matthijnssens et al, 2011; Rahman et al, 2010]. In other to answer some of the questions arising from the rapid spread of RVA genotype G12 in Africa, we investigated the Bayesian phylogeny and evolutionary dynamics of RVA genotype G12 strains in Africa.

## 2. Methods

### 2.1. Dataset

A total of 96, partial VP7 genotype G12 gene sequences spanning a period of 1998 to 2013 from twelve African countries (8 Nigerian, 11 Cameroonian, 4 Togolese, 12 Democratic republic of Congo, 3 Kenya, 12 Mozambique, 2 Ethiopia, 3 Malawi, 4 Ugandan, 17 Burkina Fasso, 1 Egypt, 1 Mauritius), were downloaded from GenBank with the help of the Rotavirus resource database. Other information collected were the date and location of isolation, any sequence without these information was not included in the dataset. Along with these were seven reference VP7 rotavirus genotype G12 genome sequences from Argentina, Thailand, Korea, and Japan. The total number of sequences in the dataset was 103. Duplicate sequences with identical year and location were excluded from the dataset.

### 2.2. Phylogenetic analysis

Sequence data was edited and assembled using CLUSTALW software (http://www.ebi.ac.uk/clustalw/). Phylogenetic trees were constructed in Mega version 6.0 software using Maximum likelihood and Neighbor joining methods with p distance model and 1000 bootstrap replicates www.megasoftware.net.

### 2.3. Evolutionary rate, time scaled Phylogeny and Population dynamic analysis

Evolutionary and additional phylogenetic analysis was carried out on African Rotavirus genotype G12 strains, using a Bayesian evolutionary approach using Markov Chain Monte Carlo (MCMC) implemented in BEAST version 1.10.4 [Drummond and Rambout, 2007]. For the MCMC run, a total of 103 rotavirus virus G12 sequences were aligned, consisting 96 African sequences (8 Nigerian, 11 Cameroonian, 4 Togolese, 12 Democratic republic of Congo, 3 Kenya, 12 Mozambique, 2 Ethiopia, 3 Malawi, 4 Ugandan, 17 Burkina Fasso, 1 Egypt, 1 Mauritius) also included are 2 reference sequences from Argentina, 2 from Japan one eachfrom Thailand and Korea, and one Porcine G12 strain. A list of the sequences used for the analysis is contained in supplementary table 1. Several models with different priors were initially evaluated strict and relaxed molecular clock models, constant population size, exponential and two non-parametric models of population changes, Bayesian Skyride plot and Gausian Markov Random Field Skyride plot. Each selected model was run for an initial 30,000,000 states. Models were then compare using the Bayes factor with marginal likelihood estimated using the path sampling and stepping stone methods implemented in BEAST version 1.10.4 [Drummond and Rambout, 2007]. Further analysis was then done using the relaxed clock with Bayesian Skyride plot coalescent prior. The MCMC run was set at 100,000,000 states with a 10% burn in. Results were visualised in Tracer version 1.8. (http://tree.bio.ed.ac.uk/software/tracer/). The effective sampling size (ESS) was calculated for each parameter, all ESS values were > 200 indicating sufficient sampling. Bayesian skyride analysis was carried out to visualise the epidemic evolutionary history using Tracer vs 1.8. (http://tree.bio.ed.ac.uk/software/tracer/). The maximum clade credibility tree was selected from the posterior tree distribution after 10% burn-in using TreeAnnotatorvs 1.8. (http://beast.bio.ed.ac.uk/TreeAnnotator/) and a time scaled Maximum scale credibility (MCC) tree was visualized in FigTree vs 1.4.

### 2.4. Phylogeographical analysis

Geographical coordinates of the locations of the African rotavirus G12 sequences were retrieved from the web with the help of online servers. A Phylogeographic tree with discrete traits was constructed using the African rotavirus G12 sequences and their geographic coordinates in latitude and longitude using a Bayesian stochastic search variable selection (BSSVS) model implemented in BEAST[Drummond and Rambout, 2007]. The discrete trait model allows for incorporation of ecological data with evolutionary analysis, it also acts as a probabilistic substitution model between discrete categories (for instance in our analysis host or location) [Drummond et al, 2002; Beale et al, 2016]. The clock prior used for the African G12 sequences were the same as the one used for the population demography. The resulting tree was annotated in Treeannotator after discarding a 10% burn-in, and visualised in Figtree. The resulting tree was spatially projected and converted to a Java script object file (JSON) and rendered into Data Driven Document (D3) library using the software SPREAD3 [Bielejec et al, 2016] after which it was visualised in HTML format using the SPREAD3 software.

## 3. Results and Discussion

The current study reports the evolutionary dynamics and phylogeoraphy of rotavirus genotype G12 in Africa. Rotavirus genotype G12 has been reported globally, it is reported that a single lineage of this genotype is responsible for its rapid global spread [Matthijnssens et al, 2010]. We analysed, 96 partial rotavirus genotype G12VP7 sequences from 13 African countries along with 7 global reference sequences. Phylogenetic analysis of the sequences revealed that all the African sequences fell into lineage III diversifying into two major clades, the West African clade highlighted in sky blue, and the East/South African clade highlighted in grey. Lineage 2 reference isolates are indicated in Red, while the lineage IV Porcine isolate is indicated in bright green (Figure 1). Our results show the African rotavirus G12 isolates have evolved into two sub lineages defined largely by geographical location. The reason for this genetic diversification has not been well defined, although factors such as natural boundaries to mass migration of human population such as the high mountain ranges of Central and East Africa, as well as the Sahara desert could serve as major factors. A similar observation of geographically bound genetic diversification was reported in a recent study of Lassa fever virus in Nigeria 2018 [Siddle et al, 2018], where genetic diversification was restricted by rivers which acted as barriers to the cross migration of rodent reservoirs of the virus.

**Figure 1.**
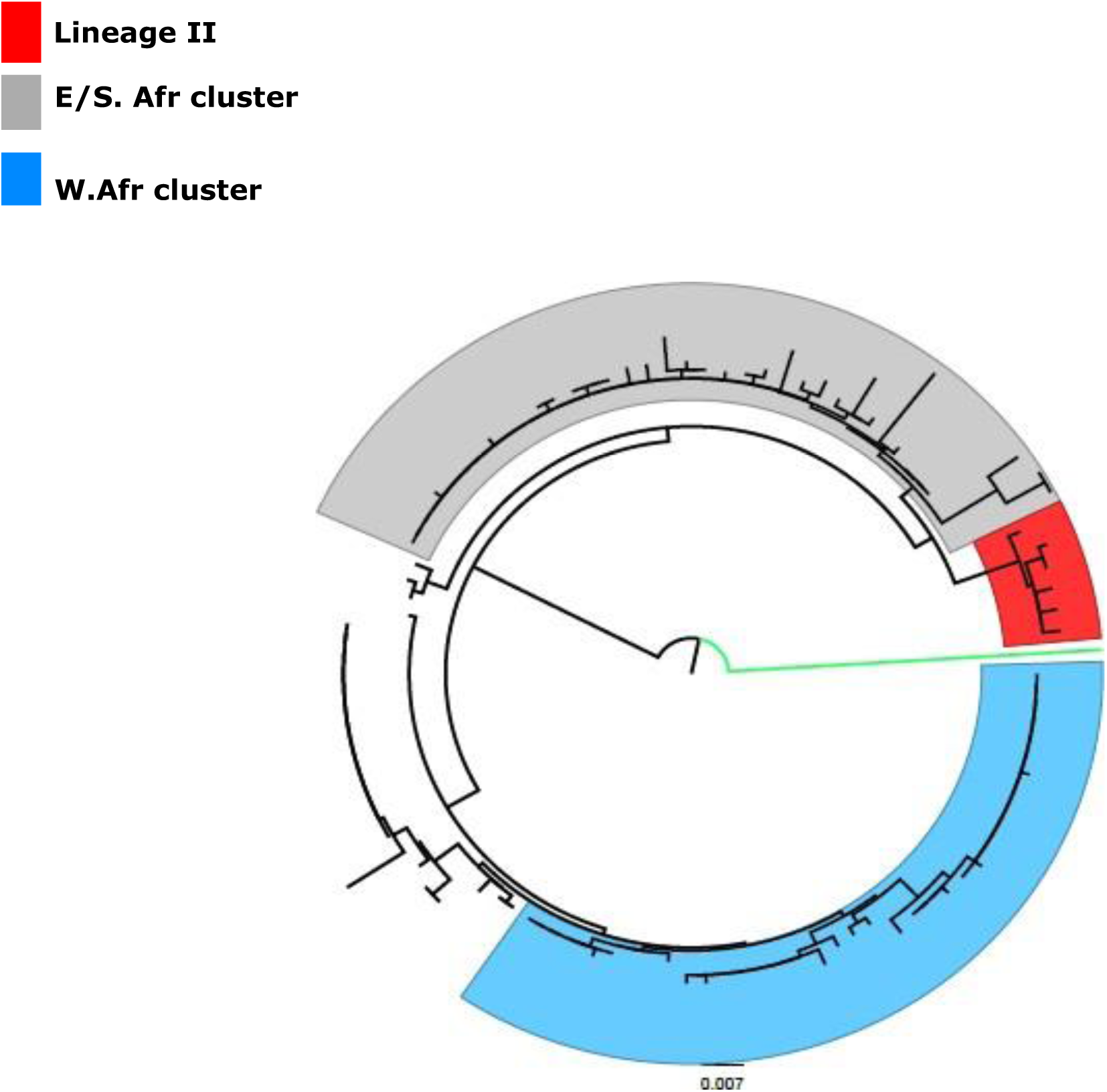
Phylogenetic tree of Rotavirus genotype G12 partial VP7 gene sequences. Tree was constructed using the Neighbour joining algorithm, with 1000 bootstrap replicates using MEGA version 6.0 software and visualised in Figtree. The West African cluster is highlighted in sky blue, the East/South African cluster highlighted in grey. Lineage 2 reference isolates are indicated in Red, while the lineage IV Porcine isolate is indicated in bright green. Scale bar is indicates number of substitutions per site.

Majority of RNA viruses have been reported to have an evolutionary rate of between 1.0×10^-3^ to 1.0×10^-6^ subs/site/year [Jenkins et al, 2002]. The evolutionary rate of the analysed African rotavirus G12 sequences was estimated to be 1,678×10^-3^, with 95% high posterior density interval (HPD) of 1.201×10^-3^ to 2.198×10^-3^ nucleotide substitutions/ site/ year (subs/site/year). This is slightly lower than the evolutionary rate of 1.89×10^-3^ of rotavirus genotype G9 reported in a global study of rotavirus molecular evolution in 2010 [Matthijnssens et al, 2010]. The evolutionary rate observed in our study hovers around the global rate previously reported for rotavirus genotype G12 in 2010 [Matthijnssens et al, 2010], this shows that the African strains have a steady mutation rate but possess the potential to diversify rapidly through molecular evolution. The time to most recent common ancestor TMRC, was calculated to be 16.8, dating back to around mid-1996. The time scaled MCC tree topology clustered into three lineages (II, III, IV), with all the African trains falling into lineage III. The African G12 strains further diversified into two main sub-clades within lineage III, the West African and South African clusters as shown in figure 2. The porcine G12 isolate was the only lineage IV strain, while the reference strains from Argentina, Thailand and Korea fell into lineage II. From the MCC tree the diversification of the 3 clusters within the African lineage III isolates occurred around the same time, between the year 2003 and 2004. The MCC tree topology follows a similar trend with the neighbour phylogeny shown in Figure 1, further buttressing the earlier mentioned observation of geographically bound diversification of African rotavirus genotype G12 VP7 genes.

**Figure 2.**
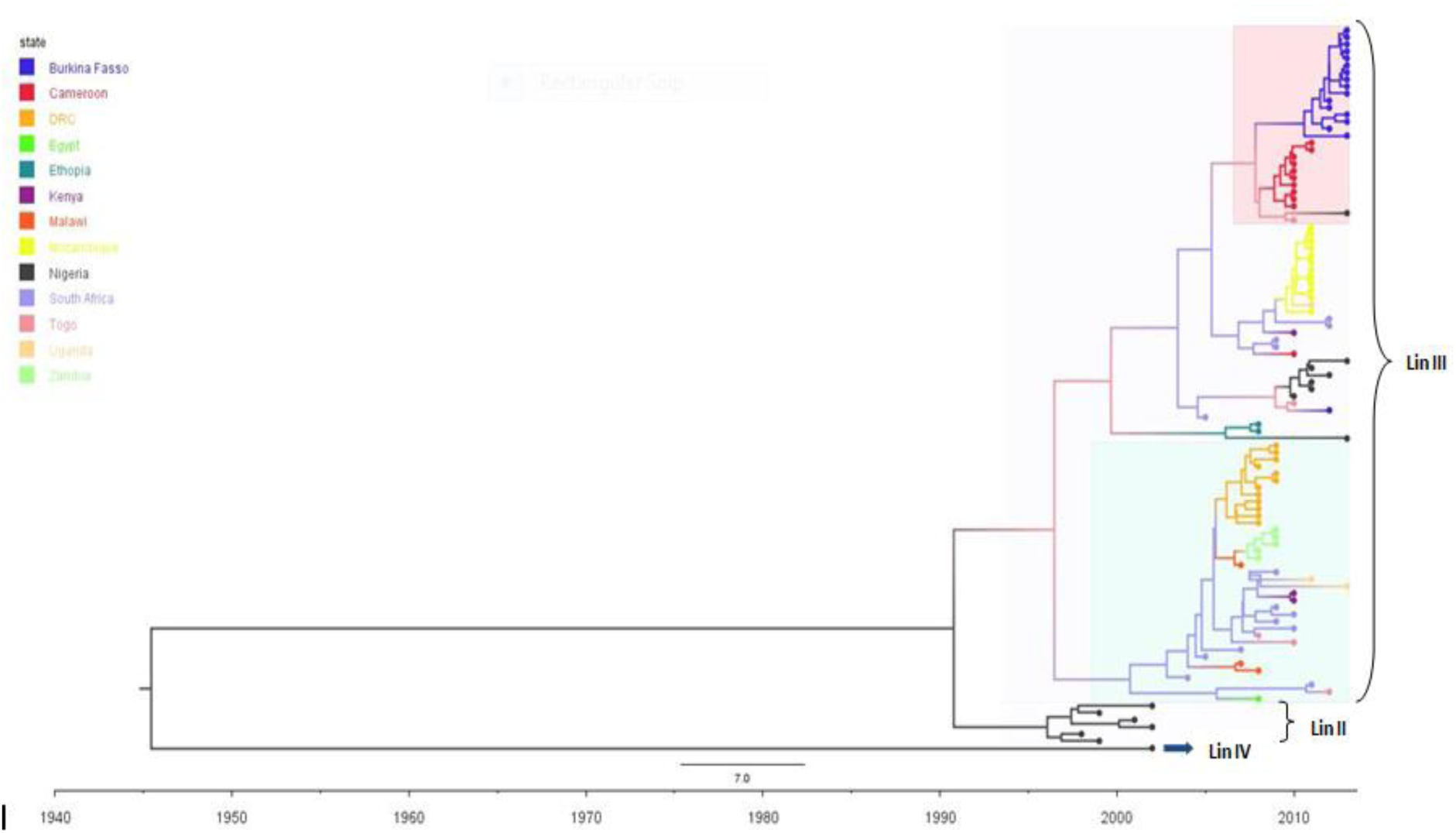
Time scaled Bayesian MCC tree of African Rotavirus virus VP7 genotype G12 sequences. Nigerian strains are indicated in Black, along with the reference G12 strains from Korea, Japan and the Porcine strain. Burkina Fasso strains are indicated in Navy blue, Cameroonian strains are indicated in red, Democratic Republic of Congo are indicated in Orange, Zambian strains are in Light green. South African strains are indicated in Sky blue, Mozambique are indicated in Yellow, Kenyan strains are indicated in Purple. Ugandan strains are indicated in Light orange, while Togolese strains are indicated in Light purple. The reference porcine strain is indicated by blue arrow. The faint Pink horizontal box represents the West African clade, while the faint Blue horizontal box represents the South African clade, within lineage III. Lineages are indicated by horizontal brackets.

The phylogeographical analysis indicated Nigeria as the most likely country of origin of rotavirus genotype G12 in Africa. The genotype then spread from Nigeria, across to South Africa (Figure 3), before moving northward to Malawi, Ethiopia and Egypt. A detailed video of the phylogeographic dispersal can be assessed in supplementary video 1. From our analysis rotavirus G12 was rapidly dispersed between countries covering long distances within a short period of time. Previous studies have shown the contribution of international travel driven by trade and economic migration to the dispersal of infectious agents across long distance countries [Zeller et al, 2015]. Asides from the long distance dispersal, the genotype was also dispersed from Nigeria, across to neighbouring West African countries of Togo, Cameroon and Burkina Fasso, showing that rotavirus genotypes are capable of both short distance and long distance spread. South Africa was identified to play a major role in the dispersal of the genotype to other regions of the continent. This might not portray the actual picture because of poor molecular surveillance in most parts of Africa [Seheri et al, 2014], but gives a clue to the viral dispersal events within the time frame (1998 to 2013). However there have been reports of several rotavirus G12 outbreaks from southern Africa [Seheri et al, 2014; Nakagomi et al, 2017; Strydom et al, 2019]. A major limitation to our study was the limited number of African rotavirus genotype G12 sequences available in GenBank during the time frame of data collection (1998 to 2013).

**Figure 3.**
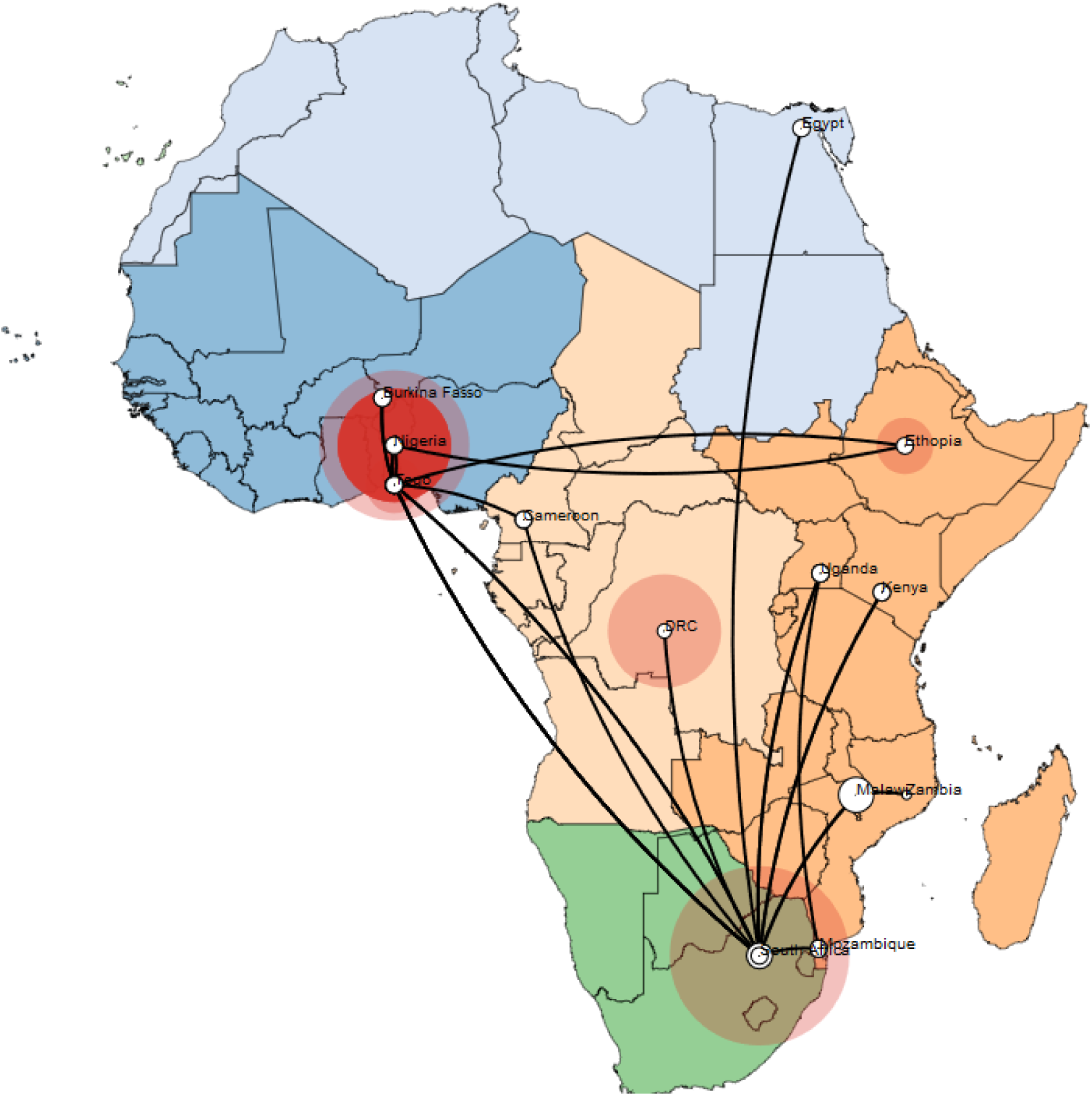
Map showing spatiotemporal viral diffusion of rotavirus genotype G12 across Africa. The map is coloured according to geographical regions. The size of the circles in each country is proportional to the number of sequences analysed for that country.

The demographic history of the African G12 viruses estimated through the BSP model, suggested that the genotype experienced a constant population dynamic (constant effective number of infections) there was a slight increase in its population size between 2004 and 2008 (Figure 4).

**Figure 4.**
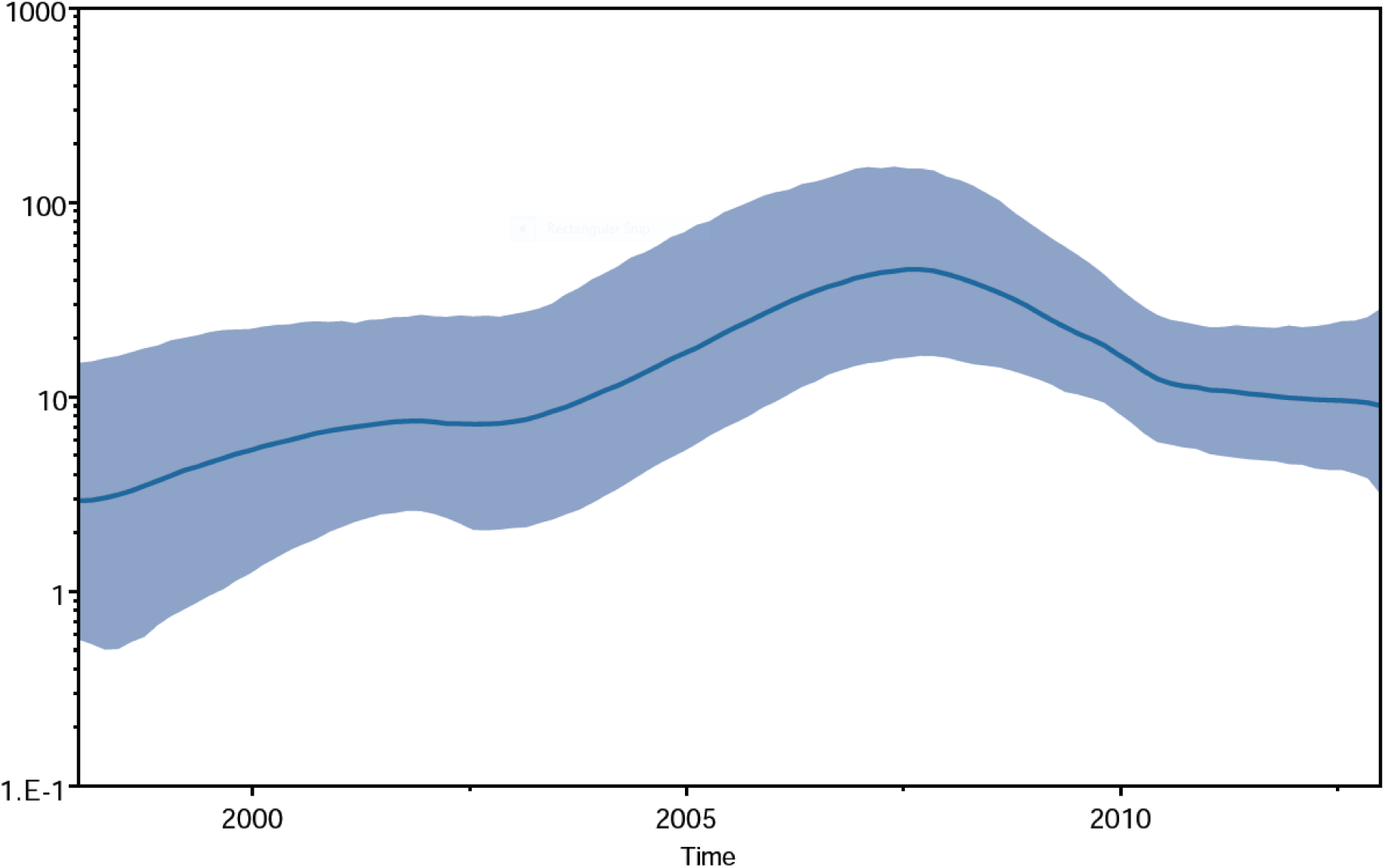
Bayesian skyplot reconstruction of African Rotavirus genotype G12strains, showing the median exponential growth line, with the blue solid area representing the 95% HPD for the growth.

## 4. Conclusions

We have shown that Rotavirus genotype G12 has diversified based on geographical locations. There is also tendency for further diversification due to its evolutionary rate and fast spread. However the population dynamic of the genotype in Africa seems to be gradually declining. This shows the positive impact of Universal routine vaccination which has been implemented in some African countries, and is being advocated for in others. We recommend the adoption of Molecular surveillance across Africa to further control spread and diversification of Rotavirus.

## Supplementary materials description

Supplementary table list the names and ascension numbers of the sequences downloaded from Genbank utilised in the analysis of the data in this study.

