## Supplementary material for "Phylogeography and Evolutionary analysis of African Rotavirus A genotype G12 reveals district genetic diversification within Lineage III": Supplimentary table 1 (from another version of this preprint: version 1, 2019-03-28)

Supplementary table 1: African group A rotavirus VP7 genotype G12 sequences downloaded from GenBank, including global reference sequences

| S/No | Isolate name | Country | Year | Ascension no |
| --- | --- | --- | --- | --- |
| 1 | RVA/R24/2010/G12P[8] | Nigeria | 2010 | JN871678.1 |
| 2 | RVA/R30/2011/G12P[8] | Nigeria | 2011 | JN871679.1 |
| 3 | RVA/R27/2011/G12P[8] | Nigeria | 2011 | JN871680.1 |
| 4 | RVA/R35/2011/G12P[8] | Nigeria | 2011 | JN871681.1 |
| 5 | RVA/TGO/MRC-DPRU5144/G12P[8] | Togo | 2010 | KJ751596.1 |
| 6 | RVA/TGO/MRC-DPRU1689/G12P[6] | Togo | 2008 | KJ752388.1 |
| 7 | RVA/TGO/MRC-DPRU5171/G12P[6] | Togo | 2010 | KJ752601.1 |
| 8 | RVA/NGR/37-2/G12P[4] | Nigeria | 2013 | KM245612.1 |
| 9 | RVA/NGR/38-2/G12P[8] | Nigeria | 2013 | KM245613.1 |
| 10 | RVA/CMR/MA01/G12P[8] | Cameroon | 2010 | KM660390.1 |
| 11 | RVA/CMR/MA11/G12P[8] | Cameroon | 2010 | KM660391.1 |
| 12 | RVA/CMR/MA24/G12P[8] | Cameroon | 2010 | KM660392.1 |
| 13 | RVA/CMR/MA46/G12P[8] | Cameroon | 2010 | KM660394.1 |
| 14 | RVA/CMR/MA70/G12P[8] | Cameroon | 2010 | KM660396.1 |
| 15 | RVA/CMR/ES283/G12P[6] | Cameroon | 2010 | KM660398.1 |
| 16 | RVA/CMR/MA88/G12P[8] | Cameroon | 2011 | KM660399.1 |
| 17 | RVA/CMR/BA356/G12P[6] | Cameroon | 2010 | KM660405.1 |
| 18 | RVA/CMR/MA02/G12P[8] | Cameroon | 2010 | KM660415.1 |
| 19 | RVA/CMR/MA60/G12P[8] | Cameroon | 2010 | KM660419.1 |
| 20 | RVA/CMR/MA127/G12P[8] | Cameroon | 2011 | KM660420.1 |
| 21 | RVA/MUS/DPRU308/2012 |  | 2012 | KP752806.1 |
| 22 | RVA/TGO/DRU4578/2010 | Togo | 2010 | KP752950.1 |
| 23 | RVA/Nig/14G1037/G12P[8] | Nigeria | 2012 | KT952028.1 |
| 24 | RVA/Nig/14G1039/G12P[8] | Nigeria | 2012 | KT952030.1 |
| 25 | RVA/BFA/2/G12P[8] | Burkina Fasso | 2012 | MG602291.1 |
| 26 | RVA/BFA/5/G12P[8] | Burkina Fasso | 2012 | MG602292.1 |
| 27 | RVA/BFA/6/G12P[8] | Burkina Fasso | 2012 | MG602293.1 |
| 28 | RVA/BFA/7/G12P[8] | Burkina Fasso | 2012 | MG602294.1 |
| 29 | RVA/BFA/14/G12P[8] | Burkina Fasso | 2013 | MG602299.1 |
| 30 | RVA/BFA/15/G12P[8] | Burkina Fasso | 2013 | MG602300.1 |
| 31 | RVA/BFA/17/G12P[8] | Burkina Fasso | 2013 | MG602301.1 |
| 32 | RVA/BFA/18/G12P[8] | Burkina Fasso | 2013 | MG602302.1 |
| 33 | RVA/BFA/19/G12P[8] | Burkina Fasso | 2013 | MG602303.1 |
| 34 | RVA/BFA/22/G12P[8] | Burkina Fasso | 2013 | MG602305.1 |
| 35 | RVA/BFA/24/G12P[8] | Burkina Fasso | 2013 | MG602307.1 |
| 36 | RVA/BFA/25/G12P[8] | Burkina Fasso | 2013 | MG602308.1 |
| 37 | RVA/BFA/27/G12P[8] | Burkina Fasso | 2013 | MG602309.1 |
| 38 | RVA/BFA/34/G12P[8] | Burkina Fasso | 2013 | MG602312.1 |
| 39 | RVA/BFA/35/G12P[8] | Burkina Fasso | 2013 | MG602313.1 |
| 40 | RVA/BFA/38/G12P[8] | Burkina Fasso | 2013 | MG602316.1 |
| 41 | RVA/COD/KisB323/G12P[6] | Congo | 2008 | KJ870790.1 |
| 42 | RVA/COD/KisB328/G12P[6] | Congo | 2009 | KJ870794.1 |
| 43 | RVA/COD/KisB508/G12P[6] | Congo | 2009 | KJ870804.1 |
| 44 | RVA/COD/KisB511/G12P[6] | Congo | 2009 | KJ870811.1 |
| 45 | RVA/COD/KisB526/G12P[6] | Congo | 2009 | KJ870824.1 |
| 46 | RVA/COD/KisB534/G12P[6] | Congo | 2009 | KJ870831.1 |
| 47 | RVA/COD/KisB539/G12P[6] | Congo | 2009 | KJ870836.1 |
| 48 | RVA/COD/KisB608/G12P[6] | Congo | 2008 | KJ870869.1 |
| 49 | RVA/COD/KisB609/G12P[6] | Congo | 2008 | KJ870870.1 |
| 50 | RVA/COD/KisB610/G12P[6] | Congo | 2008 | KJ870871.1 |
| 51 | RVA/COD/KisB613/G12P[6] | Congo | 2008 | KJ870874.1 |
| 52 | RVA/COD/KisB621/G12P[6] | Congo | 2008 | KJ870916.1 |
| 53 | RVA/EGY/AS1002/G12P[6] | Egypt | 2012 | KX265696.1 |
| 54 | RVA/ETH/DPRU5002/G12P[8] | Ethiopia | 2010 | KJ752009.1 |
| 55 | RVA/ETH/DPRU5010/G12P[8] | Ethiopia | 2010 | KJ752634.1 |
| 56 | RVA/KEN/KDH633/G12P[6] | Kenya | 2010 | AB861950.1 |
| 57 | RVA/KEN/KDH651/G12P[6] | Kenya | 2010 | AB861961.1 |
| 58 | RVA/KEN/KDH684/G12P[6] | Kenya | 2010 | AB861972.1 |
| 59 | RVA/MWI/MAL39/G12P[6] | Malawi | 2007 | AB938240.1 |
| 60 | RVA/MWI/MAL40/G12P[6] | Malawi | 2007 | AB938241.1 |
| 61 | RVA/MWI/MAL65/G12P[6] | Malawi | 2008 | AB938243.1 |
| 62 | RVA/MOZ/21125/G12P[8] | Mozambique | 2011 | KP222810.1 |
| 63 | RVA/MOZ/21134/G12P[8] | Mozambique | 2011 | KP222813.1 |
| 64 | RVA/MOZ/21136/G12P[8] | Mozambique | 2011 | KP222815.1 |
| 65 | RVA/MOZ/21140/G12P[8] | Mozambique | 2011 | KP222818.1 |
| 66 | RVA/MOZ/21141/G12P[8] | Mozambique | 2011 | KP222819.1 |
| 67 | RVA/MOZ/21144/G12P[8] | Mozambique | 2011 | KP222822.1 |
| 68 | RVA/MOZ/21186/G12P[8] | Mozambique | 2011 | KP222844.1 |
| 69 | RVA/MOZ/21188/G12P[8] | Mozambique | 2011 | KP222846.1 |
| 70 | RVA/MOZ/21193/G12P[8] | Mozambique | 2011 | KP222848.1 |
| 71 | RVA/MOZ/21194/G12P[8] | Mozambique | 2011 | KP222849.1 |
| 72 | RVA/MOZ/21195/G12P[8] | Mozambique | 2011 | KP222850.1 |
| 73 | RVA/MOZ/21206/G12P[8] | Mozambique | 2011 | KP222852.1 |
| 74 | RVA/UGA/DPRU-4616/G12P[6] | Uganda | 2011 | KJ753286.1 |
| 75 | RVA/UGA/DPRU-4620/G12P[6] | Uganda | 2011 | KJ753732.1 |
| 76 | RVA/UGA/MUL-13-183/G12P[6] | Uganda | 2013 | KX632270.1 |
| 77 | RVA/ZAF/3133WC/G12P[4] | South Africa | 2009 | HQ657154.1 |
| 78 | RVA/ZAF/DPRU5156/G12P[8] | South Africa |  | KJ751784.1 |
| 79 | RVA/ZAF/DPRU2130/G12P[8] | South Africa | 2005 | KJ751998.1 |
| 80 | RVA/ZAF/DPRU1191/G12P[8] | South Africa | 2009 | KJ752344.1 |
| 81 | RVA/ZAF/DPRU75/G12P[8] | South Africa | 2012 | KJ752366.1 |
| 82 | RVA/ZAF/DPRU1911/G12P[6] | South Africa | 2007 | KJ752474.1 |
| 83 | RVA/ZAF/DPRU4090/G12P[6] | South Africa | 2011 | KJ752819.1 |
| 84 | RVA/ZAF/DPRU138/G12P[8] | South Africa | 2009 | KJ753024.1 |
| 85 | RVA/ZAF/DPRU108/G12P[8] | South Africa | 2009 | KJ753340.1 |
| 86 | RVA/ZAF/DPRU1554/G12P[8] | South Africa | 2010 | KJ753697.1 |
| 87 | RVA/ZAF/DPRU1656/G12P[6] | South Africa |  | KJ752687.1 |
| 88 | RVA/ZAF/DPRU1370/G12P[6] | South Africa | 2004 | KJ752939.1 |
| 89 | RVA/ZAF/DPRU76/G12P[8] | South Africa | 2012 | KJ752939.1 |
| 90 | RVA/ZMB/DPRU3491/G12P[6] | Zambia | 2009 | KF636151.1 |
| 91 | RVA/ZMB/DPRU1680/G12P[6] | Zambia | 2008 | KJ752535.1 |
| 92 | RVA/ZMB/DPRU3488/G12P[6] | Zambia | 2009 | KJ752645.1 |
| 93 | RVA/ZMB/DPRU1660/G12P[6] | Zambia | 2008 | KP752871.1 |
| 94 | RVA/ZMB/DPRU3506/G12P[6] | Zambia | 2009 | KP753206.1 |
